# Changing Patterns and Drivers of Increased Pesticides Use in Smallholder Vegetable Production Systems in Tanzania

**DOI:** 10.1101/2021.01.18.427098

**Authors:** Jones A. Kapeleka, Elingarami Sauli, Omowunmi Sadik, Patrick A. Ndakidemi

## Abstract

Pesticides use has become a key component of smallholder horticulture production. Therefore, dynamics in pesticides handling need to be ascertained. This study assessed drivers of pesticides use and determinants of changing patterns of pesticides handling practices in smallholder vegetable systems. Data were collected from 385 farmers from Iringa, Arusha, Manyara, and Kilimanjaro regions in Tanzania through an in-depth survey and field observations. A binary probit model was used to derive factors fostering increased pesticides use. Results showed that 47.9 % of all pesticides were wrongly used. Most farmers (88.6%) lacked knowledge of pest control and 88.9% were unaware of safety practices. Disposal methods of empty pesticides foster occupational and environmental exposure (58%). There was an increasing trend in pesticides use (58.4%), accompanied by changing pesticides formulations. Over 60 pesticides with 29 different formulations were used. Mixing pesticides (71.2%), high dose rates with increased frequency of application were observed. Pesticides under Class II WHO hazard (68.9%) dominated. Extremely hazardous (Class Ia) and highly hazardous (Class Ib) were also used. Binary probit model showed that number of crops grown, pesticides mixing, and region contributed positively to the likelihood of increased pesticides use while farmers’ perception of effectiveness of pesticides, lack of access to safe use information, poor use of safety gears and inability to read pesticides labels had a negative impact. The fate of pesticides use in smallholder vegetable production systems is therefore the culmination of serious health and environmental implications. Excessive pesticides use escalated by increased number of crops, improper use of PPE, and pesticides mixing practices subjects the general population to pesticides environmental exposure thereby jeopardizing sustainability of smallholder vegetable production in Tanzania. Regular training to farmers and extension officers on current and emerging issues on pests and pesticide safe use is vital.

## 1. Introduction

Pesticides are extensively used in agricultural production to prevent and control insect pests, diseases, weeds, and other plant pathogens to reduce or eliminate yield losses and get high-quality product [1,2]. Their use has increased due to their rapid action. Organophosphates, carbamates, and synthetic pyrethroids are the most common pesticides in developing countries. However, these chemicals have the potential to cause adverse effects on human and environmental health [3–6].

In Tanzania, pesticides are extensively used in areas where coffee, fruits, and vegetable farming are practiced [1]. Smallholder vegetable farmers depend heavily on the use of pesticides for the control of different pests and diseases. This is probably because they believe that the only solution to pest problems is to spray more frequently and to use different types of pesticides [7]. Poor extension services, farmers who are not informed through agricultural input providers and insufficient knowledge on pesticides usage are common factors leading to overuse of different pesticides [8].

Undeniably, pesticide use will continue to be an essential component of agricultural production. This is because pesticides use produces immediate benefits to the farming population, but the long term risks are shared by society as a whole [9]. Pesticides use among smallholder farmers in Tanzania has been reported. Major areas reported include pesticides handling practices and acute poisoning resulting from exposure to pesticides [1,10–12], but there is limited information on drivers of farmers’ behaviour and determinants of multiple pesticides use across different farming typologies and their implication on the sustainability of smallholder vegetable subsector.

Compared with previous studies, this study critically assessed determinants of increased pesticides use and drivers of farmers’ behavior in changing pesticides use practices. It elucidated the causal link between pesticides use and the fate of pesticide use in smallholder vegetable production systems in Tanzania, hence providing useful information on farmers’ behavior and changing practices of pesticides use and handling practices in smallholder vegetable production systems.

## 2. Materials and Methods

### 2.1 Study area

The assessment was undertaken in highest smallholder vegetable production areas which included southern highlands and northern corridor in Tanzania. These areas are typical vegetable areas with intensive pesticide use. The study focused on smallholder vegetable crops that are widely produced and consumed in Tanzania, including tomatoes, onions, watermelons, and sweet pepper. Other leafy vegetables included Chinese cabbage, carrots, cucumbers, cabbage, amaranths, kale and night shade.

### 2.2 Sampling procedures and sample size

Purposive sampling was employed in selecting regions and districts with high vegetable productivity in different agro ecological zones. The decision to select southern highlands and northern corridor was based on the productivity of vegetable crops as well as the extensive use of pesticides. Simple random sampling was used to select wards, villages and household to be included in the study sample. A total sample of 385 farmers was used in this study. From the 420 survey questionnaires that were administered, 385correctly filled, and hence the response rate of 92%. Vegetable farmers were randomly selected from the list of households provided by respective village government officers. Randomness was achieved by assigning random numbers to the list of participants from which the sample was collected. The sample was chosen based on the proportion of farmers involved in smallholder vegetable production.

The sample size was calculated based on the following formula for determination of minimum sample size

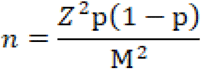

Where n = Sample size, Z = % point of the standard normal distribution which is 1.96 in this case corresponding to 95% confidence level, M = marginal error which is 5%, p= expected proportion of the respondents taken as 50%, =0.5, q = 1-p

### 2.3 Research hypothesis

Based on previous studies, this study hypothesized that in reducing the effects of the pests, farmers have resorted to using more pesticides. Demographic variables, mixing practices, poor extension services, region of the farmers, acreage, frequency of pesticides application, number of crops grown, and low general knowledge on pesticide usage were attributes to overuse of different pesticides.

### 2.4 Data collection procedures

A structured questionnaire containing both closed and open ended questions was administered to the respondents by trained interviewers. Data were collected through face-to-face interviews with the farmers and responses recorded during interviews. The questionnaire used in previous studies was used with minor modifications to suit the current research. Reliability of the questionnaire was measured by administering the improved questionnaire twice over a period of one moth to a group of farmers from Arusha region. This improved questionnaire was pretested among 20 individuals from one village in the study areas, which was finally removed from the sample.

All questionnaires were translated and administered in Kiswahili, an official language in Tanzania. Information about socio-economic variables such as age, education, pesticides application equipment, safety measures, and pesticides application technologies, landholding, access to training in pesticide use, and mixing practices were collected. Information on perception on the effectiveness of pesticides, frequency of pesticides application, types of crops grown and a number of pesticides mixed in a single mixture was also collected as explanatory variables for the determinants of farmers’ changing patterns of increased pesticides use.

### 2.5 Data Processing and Analysis

Data were analyzed using SPSS 18.0 version computer software whereby descriptive statistics such as frequencies and percentages were calculated for each variable under study and used to obtain the general picture of respondents’ profile. Information about pesticides handling, spraying techniques, and use of pesticide labels was also analyzed. Descriptive statistics, namely frequencies and means, as well as cross-tabulation was also used to summarize information among different categories of respondents. Quantitative data are presented as means and standard deviations (Mean ± SD).

Pearson correlations were used to assess the relationship and significance of the association between increased levels of pesticides use and demographic variables. Chi-squared (χ^2^) test was used to compare frequencies and to determine significant differences in the proportions of responses given. The levels of pesticides used were dichotomized into high and low at one standard deviation below the mean levels of volumes of pesticides sprayed per day. Determinants of farmers’ changing patterns of increased pesticides use were determined by regressing the levels of pesticides use (dependent variable) on a set of demographic and pesticides handling practices variables using a binary probit model. Significant level for the results was accepted at p < 0.05.

The regression model used to estimate the determinants of increased pesticides use (IPU) is given by:

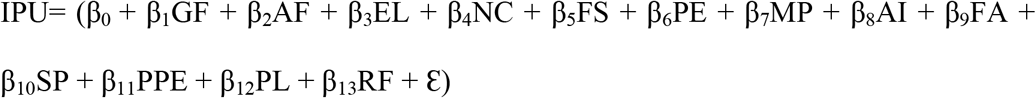

Where

IPU= Increased Pesticides Use, GF= Gender of Farmer, AF=Age of Farmer, EL= Education Level, NC= Number of vegetable Crops each farmer grows, FS= Farm Size, PE= Perception on the Effectiveness of Pesticides, MP= Mixing Practices, AI= Access to Information on Pesticide Use, FA= Frequency of Pesticides Application, SP = Source of Pesticides, PPE= Use of Personal Protection Equipment, PL=Read Pesticide Label, RF= Region of the farmer, Ԑ= unknown parameters.

The rationale of this model is to estimate the probability that observation with particular characteristics falling in one of the proposed categories [13]. Thus, this binary probit model was used to predict the likelihood that farmers will resort in increased use of pesticides based on specific predictors. The region was included in the model to ascertain possible differences in farmers’ level of pesticides use concerning their geographical location.

### Ethical clearance

Ethical clearance was obtained from Tanzania’s National Institute of Medical Research (NIMR) with Reference No. NIRM/HQ/R.8a/Vol.IX/2742. Each participants signed a written consent form for participation in the research.

## 3. Results

### 3.1 Respondent’s demographic information

Males (77.9%) dominated the smallholder vegetable production compared with females (22.1%). The farming population was relatively younger. Youths (25-34 years) and middle-aged individuals (35-44 years) were involved in smallholder vegetable production (31.7% and 33.0% respectively). The education level was generally low. Most farmers (79.4%) had attained primary level education, while only 13.5% had attained ordinary level secondary education (Table 1). Tomatoes (80.8%), onions (35%), cabbage (27.1%) and 17.1% sweet papers were the main vegetable crops grown by smallholder vegetable producers. Others included Chinese cabbage, nightshade, kale, amaranths African eggplants, cucumbers okra and carrots. The average area under production was 1.24 ± 0.81 acres (data not shown).

**Table 1.**
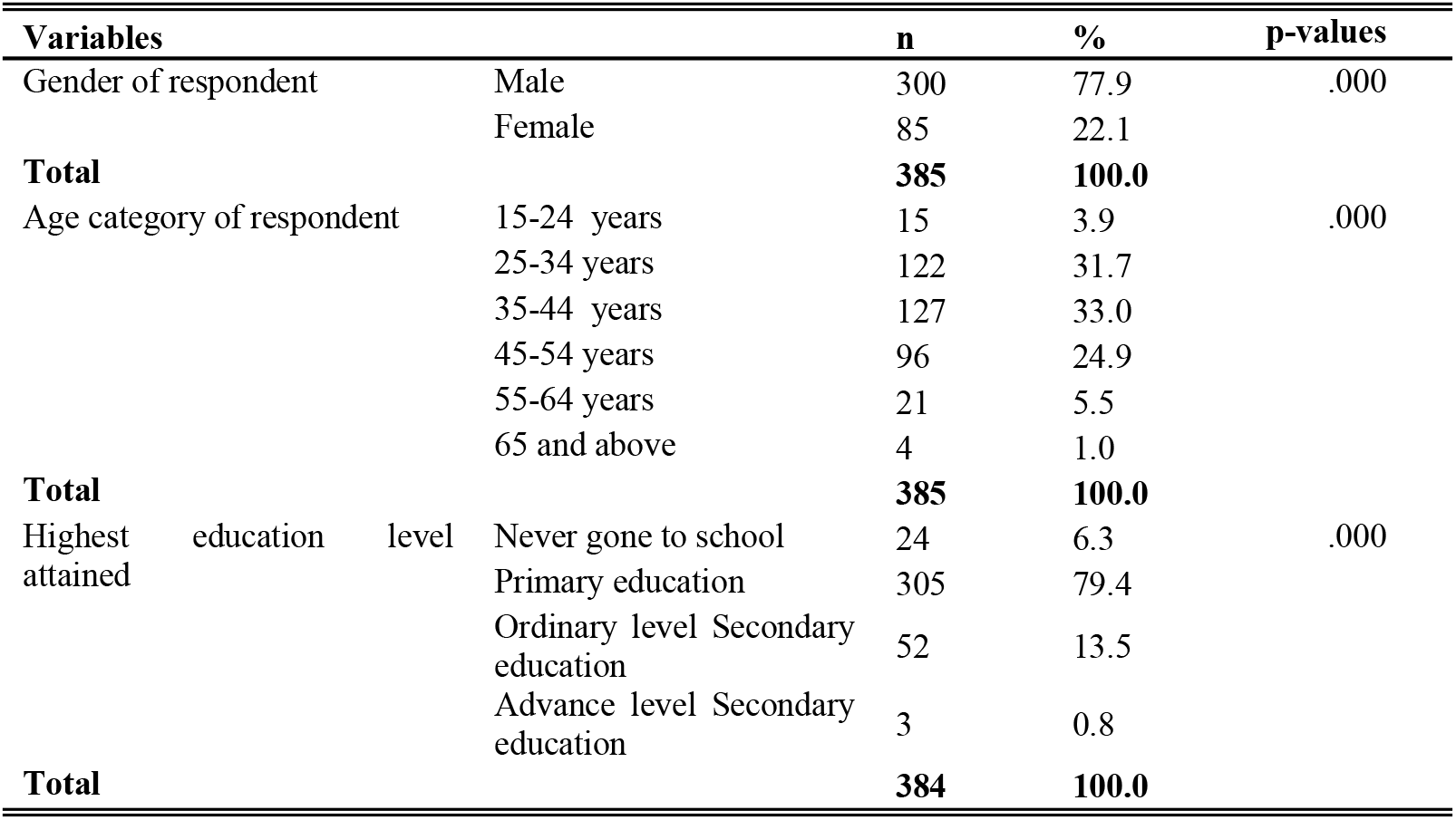
Demographic information from the study area.

### 3.2 Pesticides types and use practices

Insecticides (56.8%) and fungicides (39.2%) and to a small extent, herbicides (1.4%) were the main pesticides used. Misuse of pesticides in smallholder horticulture production was noted as well. Some farmers used acaricides for controlling vegetable pests. Of all pesticides used, 52.1% were correctly used for target crops, while farmers improperly used 47.9%. These included banned pesticides products, unregistered pesticides, and pesticides registered for use in controlling ticks (acaricides) and other crops such as coffee, cashew nuts, and ornamental flower production.

Smallholder vegetable producers have limited access to extension services to advise them on pest control. Table 2 shows that 88.6% had not received agricultural experts’ advice on pest control in the last three years. Likewise, 88.9% had not received any advice on the safe use of pesticides. Very few farmers (11.1%) had received some training on the safe use of pesticides, mainly received from pesticides retailers (47.5%), extension officers (45.0%) and NGO’s (7.5%).

**Table 2.**
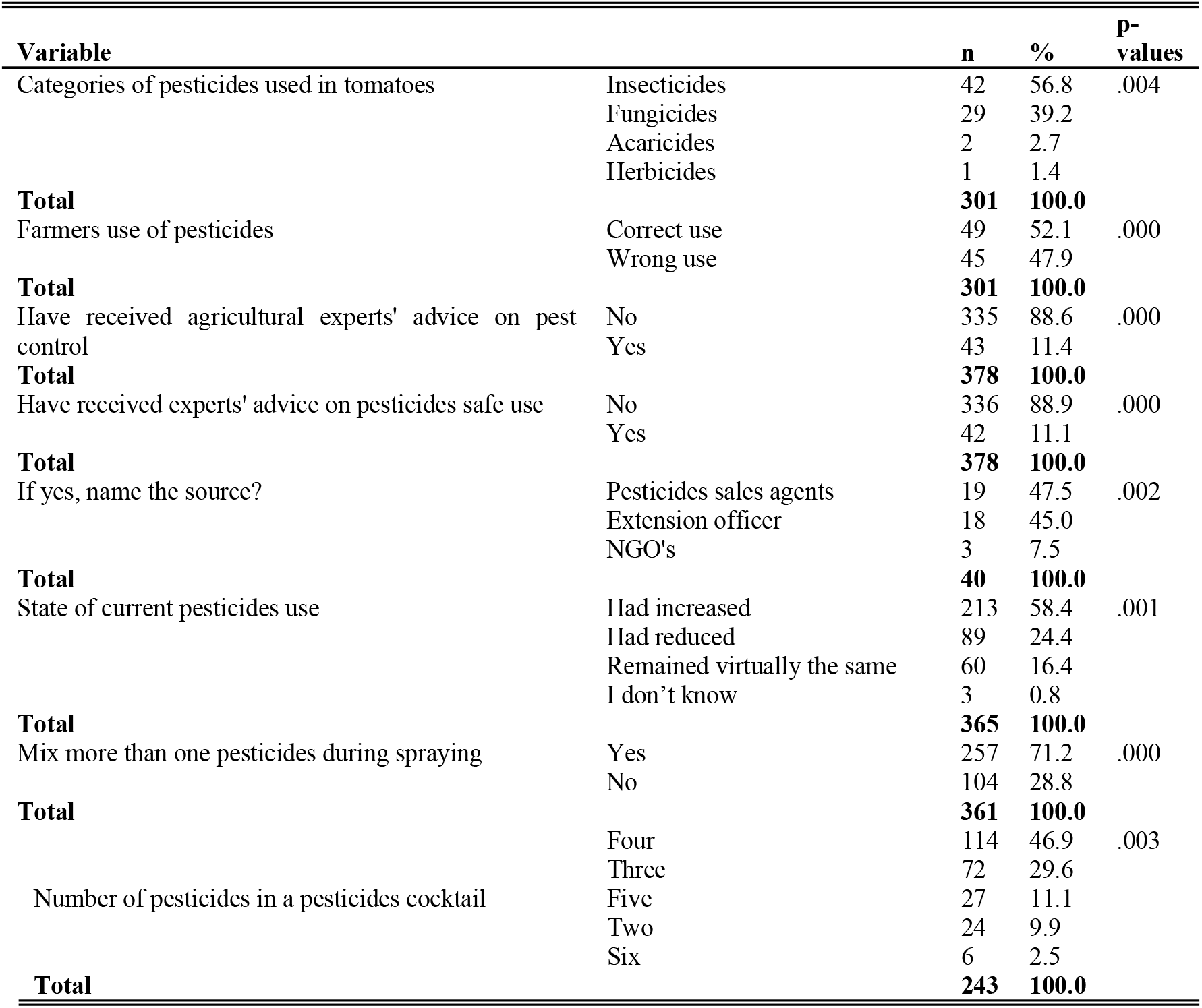
Pesticides use and practices in vegetable production.

The rate of pesticides use had increased in the recent past (58.4%). Farmers (71.2%) mixed two or more pesticides during spraying. The reasons for mixing pesticides included minimizing the spraying costs (32%), increasing pesticides effectiveness (28.1%), controlling all pests crops at once (26.1%) and ensuring that the pesticides complement each other in controlling of pests (13.8%). Up to six different pesticides were mixed in a single tank. Close to half of the farmers (46.9%) mixed four different pesticides in one mixing tank during spraying, 29.6% mixed three pesticides, while 11.1% mixed up to five different pesticides (Table 2).

### 3.3 Pesticides volumes used in vegetable production

The most used mixing equipment were drums (200 L) and knapsack sprayers (20 L) (67.8% and 32.8% respectively). High volumes of pesticides use per acre was a common practice among smallholder vegetable producers. On average, farmers spray 4.27 drums (758.7 L/acre) of pesticides mixture on an average of 1.21 acre of land in tomato, while 4.59 drums (913.75 L/acre) of pesticides mixtures are sprayed on an average of 1.08 acre of onion (Table 3).

**Table 3.** Pesticides volumes used in vegetable production.

| Variable | Equipment used in mixing |  |  |  |  |  |
| --- | --- | --- | --- | --- | --- | --- |
|  | Drum |  |  | Knapsack |  |  |
|  | Minimum | Maximum | Mean | Minimum | Maximum | Mean |
| Tomato farm size in acres | 0.25 | 5.00 | 1.21 | 0.25 | 2.00 | 0.97 |
| Number of drums/knapsacks per day | 1.00 | 12.00 | 4.27 | 1.00 | 50.00 | 14.14 |
| Spraying hours/day | 1.00 | 12.00 | 5.06 | 1.00 | 12.00 | 4.48 |
| Number of spray men | 1.00 | 10.00 | 3.33 | 1.00 | 5.00 | 2.44 |
| Number of spraying days | 1.00 | 20.00 | 2.72 | 1.00 | 3.00 | 1.04 |
| Onion farm size in acres | 0.25 | 5.00 | 1.08 | 0.25 | 1.00 | 0.55 |
| Number of drums (200l) per day | 1.00 | 16.00 | 4.59 | 1.00 | 8.00 | 3.33 |
| Spraying hours/day | 1.00 | 12.00 | 5.12 | 1.00 | 9.00 | 2.77 |
| Number of spray men | 1.00 | 13.00 | 2.71 | 1.00 | 8.00 | 1.59 |
| Number of spraying days | 1.00 | 24.00 | 1.79 | 1.00 | 1.00 | 1.00 |

### 3.4 Pesticides use and management of empty pesticides containers

The main disposal method of empty pesticides containers was burning (58%). Other farmers reported burying (15%), throwing in dustbins (12%), leaving empty containers in the fields (8%) and very few farmers use empty pesticides for storing water and food (3% and 2%), respectively. Most farmers (70.3%) did not read pesticides labels before using pesticides and 83.8% did not follow the instructions. Spraying frequency of pesticides was relatively high among the farmers. Sixty-one percent (61%) sprayed pesticides once a week, 18% sprayed twice a week, and 12% sprayed once in two weeks.

Most pesticides (78.5%) used in smallholder vegetable production had full registration status, while (19.4%) were not registered. Furthermore, 68.9% of all reported pesticides fell under Class II (Moderately Hazardous) of WHO hazard classification of pesticides, while 24.3% were under Class U (Unlikely to present acute hazard in normal use). Small quantities of extremely hazardous (Class Ia) and highly hazardous (Class Ib) were also found to be used in vegetable production by smallholder farmers. About three quarters (76.6%) placed pesticides in a pesticide store while others stored them in their farms, general store, and bathrooms/toilets (Table 4).

**Table 4.**
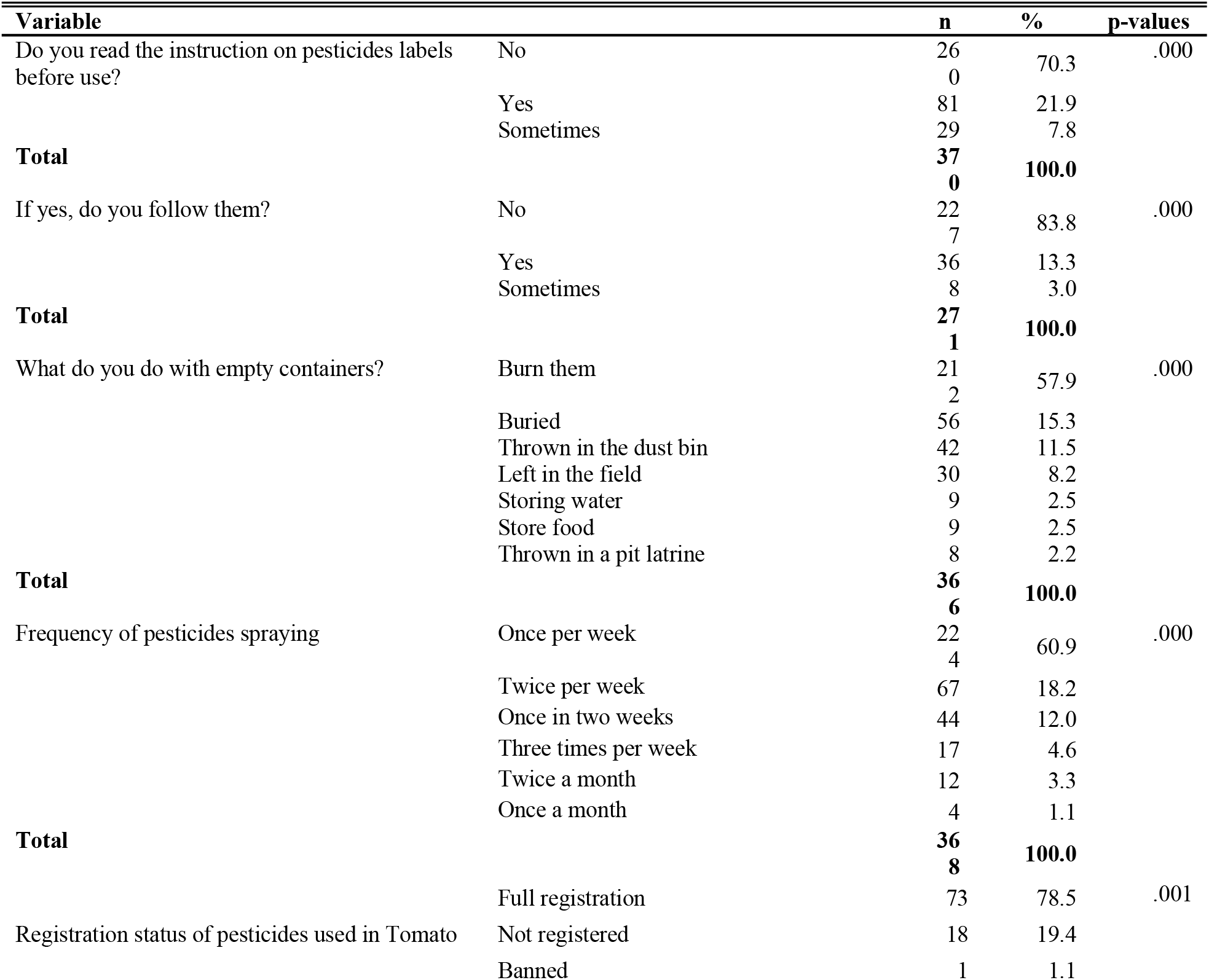
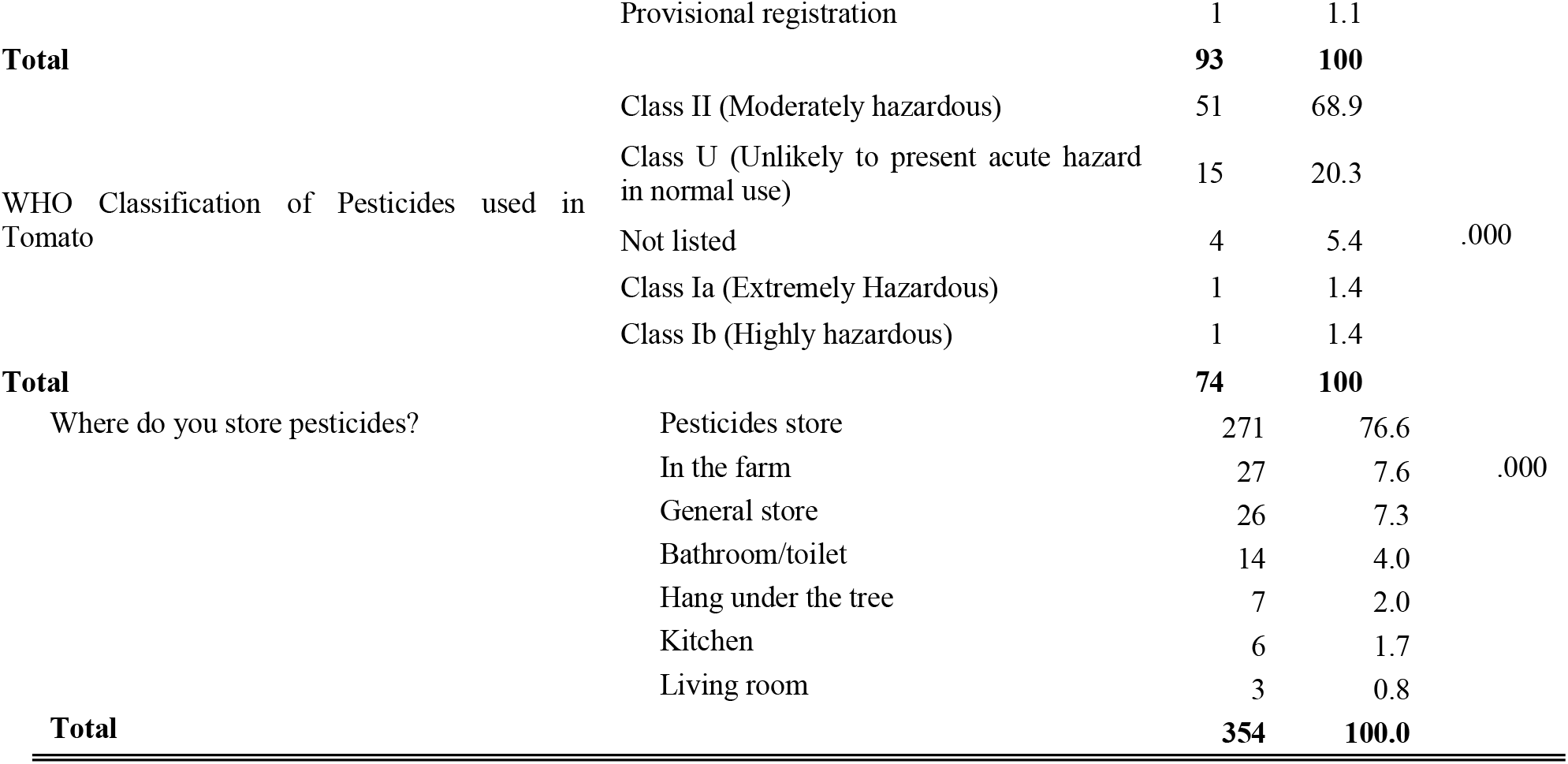
Pesticides use and management of empty pesticides containers.

Over 60 different pesticides were used in tomato production only comprised of 29 different formulations. Organophosphorus (97.6%), Carbamates (54.1%), and Substituted benzene (34.6%), a combination of Pyrethroid+Organophosphorus (28.8%), Avermectin (28.1%), a combination of Carbamate+Acylalanine (22.9%) and Dithiocarbamate (19.5%) constitute main chemical families of pesticides used (Table 5).

**Table 5.**
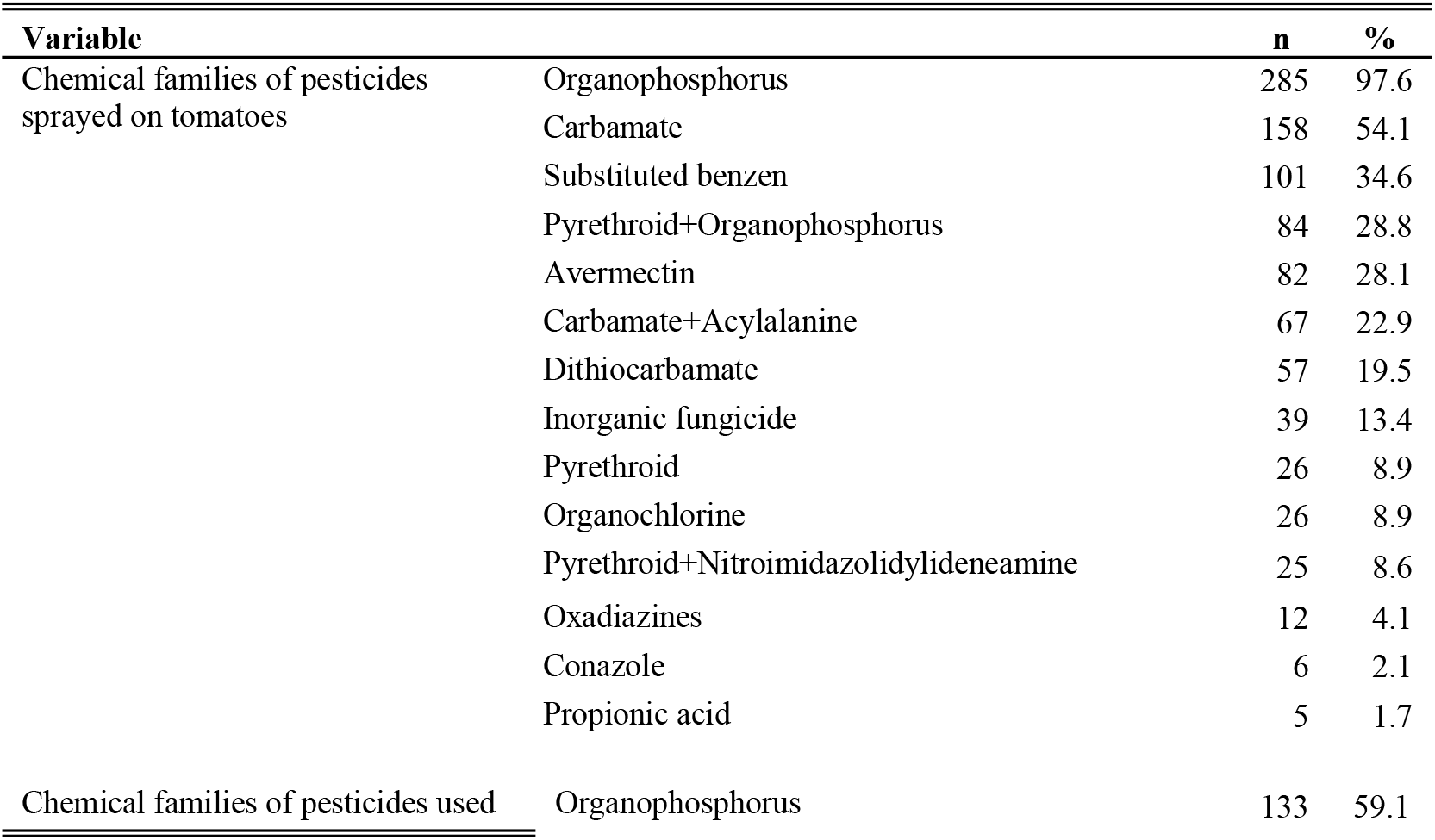
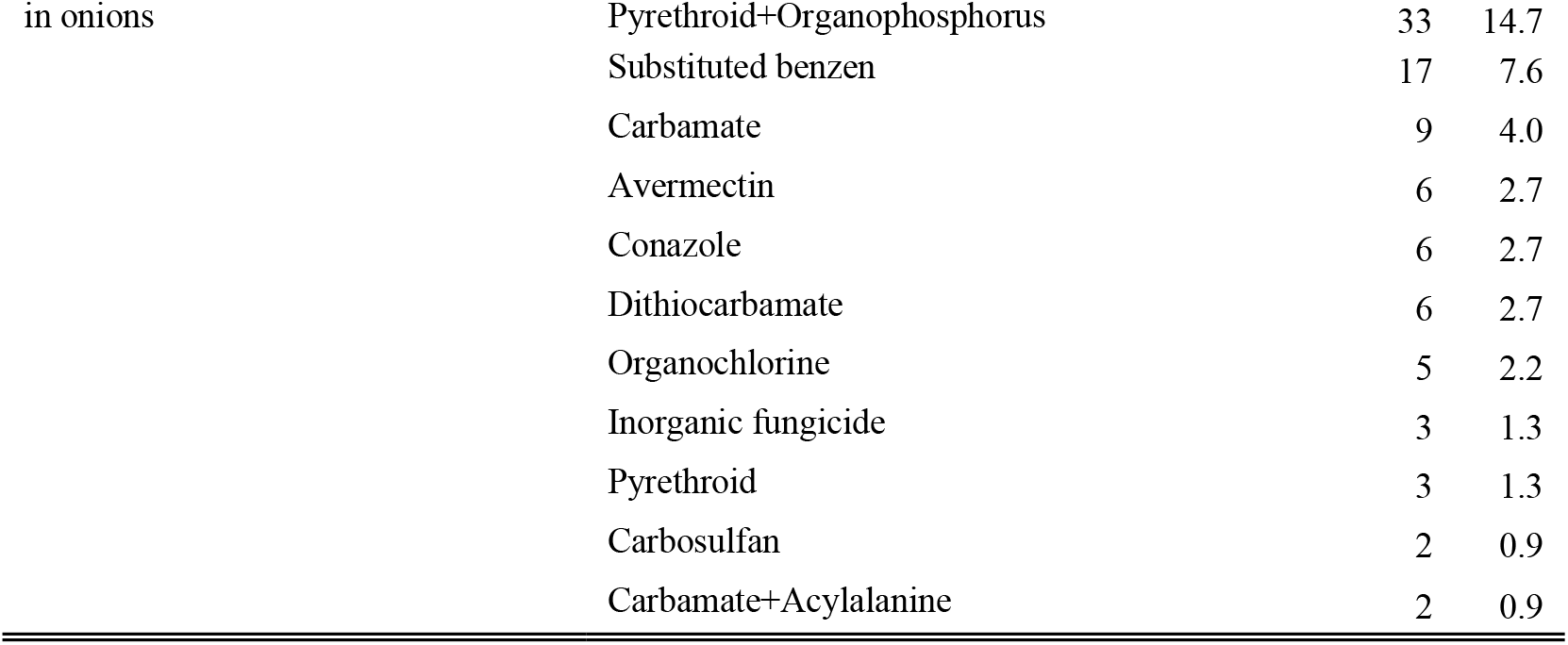
Pesticides chemical families used smallholder vegetable production.

### 3.5 Pesticides safety and use of personal protection equipment (PPE)

Farmers did not have the basic and essential personal protective equipment for use during pesticides handling. Close to three quarters (74.5%) did not wear gloves when handling pesticides, 86.6% did not use respirators, 82.6% had no masks, 91.5% did not wear goggles, and 79.3% did not wear overalls when handling pesticides. Gumboots (60.3%) were the only PPE used mostly in tomato fields, while some farmers in onion fields were observed spraying barefooted (Table 6).

**Table 6.** Personal Protective Equipment used by farmers.

| Variable |  | n | % | p-values |
| --- | --- | --- | --- | --- |
| Gloves | No | 272 | 74.5 | .000 |
|  | I use and is in good order | 73 | 20.0 |  |
|  | Have but don't use | 20 | 5.5 |  |
| **Total** |  | **365** | **100.0** |  |
| Boots | I use and is in good order | 219 | 60.3 | .000 |
|  | No | 119 | 32.8 |  |
|  | Have but won out | 21 | 5.8 |  |
|  | Have but don't use | 4 | 1.1 |  |
| **Total** |  | **363** | **100.0** |  |
| Respirator | No | 309 | 86.6 | .000 |
|  | I use and is in good order | 42 | 11.8 |  |
|  | Have but don't use | 6 | 1.7 |  |
| **Total** |  | **357** | **100.0** |  |
| Mask | No | 300 | 82.6 | .000 |
|  | I use and is in good order | 53 | 14.6 |  |
|  | Have but won out | 8 | 2.2 |  |
|  | Have but don't use | 2 | 0.6 |  |
| **Total** |  | **363** | **100.0** |  |
| Goggles | No | 332 | 91.5 | .000 |
|  | I use and is in good order | 21 | 5.8 |  |
|  | Have but don't use | 6 | 1.7 |  |
|  | Have but won out | 4 | 1.1 |  |
| **Total** |  | **363** | **100.0** |  |
| Overalls | No | 288 | 79.3 | .000 |
|  | I use and is in good order | 66 | 18.2 |  |
|  | Have but don't use | 8 | 2.2 |  |
|  | Have but won out | 1 | 0.3 |  |
| **Total** |  | **363** | **100.0** |  |
| Head cover | No | 307 | 85.0 | .000 |
|  | I use and is in good order | 48 | 13.3 |  |
|  | Have but don't use | 6 | 1.7 |  |
| **Total** |  | **361** | **100.0** |  |

### 3.6 Drivers of increased pesticide use

Generally, farmers used increased application rates with high pesticides volumes both in tomato and onion production systems. The correlation coefficients between increased pesticides use and demographic variables showed that increased pesticides use had a significant positive correlation with mixing more than one pesticide during spraying, number of crops grown consecutively, and region of the respondent. On the other hand, increased pesticides use had a significant negative correlation with access to safe use information, perception of the effectiveness of pesticides, wearing of personal protection equipment and reading the instruction on pesticides labels before use (Table 7).

**Table 7.** Correlation test between the increased use of pesticides and demographic variables.

| Variable | Pearson Correlation | p-value |
| --- | --- | --- |
| Gender of respondent | -0.119 | 0.055 |
| Age category of respondent | -0.114 | 0.065 |
| Highest education level attained | -0.009 | 0.886 |
| Access to safe use information | -0.142* | 0.022 |
| Perception on effectiveness of pesticides | -0.143* | 0.022 |
| Mix more than one pesticides during spraying | 0.225** | 0.000 |
| Number of crops grown consecutively | 0.264** | 0.000 |
| Do you wear personal protective equipment | -0.258** | 0.000 |
| Frequency of pesticides spraying | -0.001 | 0.982 |
| Region of respondent | 0.553 | 0.000 |
| Read instruction on pesticides labels before use | -0.183** | 0.003 |

\*. Correlation is significant at the 0.05 level (2-tailed).
\*\*. Correlation is significant at the 0.01 level (2-tailed).

Determinants of farmer’ increased use of pesticides are presented in table 8 from the results of the probit regression model. The predictors tested were: region of the farmer, gender, age, education level, number of vegetable crops each farmer grows, farm size, perception on the effectiveness of pesticides, mixing practices, frequency of pesticides application, access to information on pesticide use, source of pesticides, use of personal protection equipment and the tendency of farmers to read pesticide label. Among the variables tested, the region of the farmer, the number of vegetable crops grown and mixing of pesticides positively contributed to the likelihood of increased use of high levels of pesticides. On the contrary, farmers’ perception of low effectiveness of insecticides, access to information on pesticides safe use, use of safety measures, and reading pesticides labels negatively influenced increased use of the high level of pesticides. In other words, the perception of low effectiveness of pesticides, limited access to information on the safe use of pesticides, low use of protective gears among farmers increased the likelihood of farmers using pesticides indiscriminately. Other variables including gender, age category, level of education, farm size, frequency of pesticides spray, source of pesticides showed no significant contribution in influencing farmers’ decision to excessively apply high levels of pesticides.

**Table 8.** Drivers of farmers’ increased use (Results of the Binary Probit Model)

| Explanatory variables | Estimate | Std. Error | Sig. | Wald Chi-Square | 95% Wald Confidence Interval |  |
| --- | --- | --- | --- | --- | --- | --- |
|  |  |  |  |  | Lower | Upper |
| Region | 1.488 | 0.2236 | 0.000 | 44.273 | 1.049 | 1.926 |
| Gender | 0.65 | 0.2921 | 0.126 | 4.957 | 0.078 | 1.223 |
| Age category | 0.254 | 0.162 | 0.117 | 2.451 | -0.064 | 0.571 |
| Highest level of education | 0.107 | 0.2593 | 0.68 | 0.17 | -0.401 | 0.615 |
| Number of crops grown | 0.147 | 0.0468 | 0.002 | 9.919 | 0.056 | 0.239 |
| Farm size | -0.395 | 0.2077 | 0.057 | 3.616 | -0.802 | 0.012 |
| Perception on effectiveness of pesticides | -0.555 | 0.2429 | 0.022 | 5.221 | -1.031 | -0.079 |
| Mixing of pesticides | 0.592 |  | 0.012 | 6.293 | 0.129 | 1.054 |
| Access to safe use information | -0.717 | 0.4284 | 0.04 | 4.205 | -1.718 | -0.039 |
| Frequency of pesticides spray | -0.2 | 0.0932 | 0.116 | 2.47 | -0.329 | 0.036 |
| Source of pesticides | -0.07 | 0.3145 | 0.823 | 0.05 | -0.687 | 0.546 |
| Wearing of PPEs | -0.349 | 0.1698 | 0.04 | 4.222 | -0.682 | -0.016 |
| Reading of Pesticides Label | -0.446 | 0.2136 | 0.037 | 4.351 | 0.027 | 0.864 |

## 4. Discussion

Pesticides will continue to be an essential component of agricultural production in farmers’ efforts to control pests and increase farm income. In this study, determinants of increased pesticides use and drivers of farmers’ behavior in changing pesticides use practices were assessed. The novelty of this current study emanates from the establishment of causal link of increased pesticides use with mixing practices, farmers’ perception on the effectiveness of pesticides, crops each farmer grows, pesticides handling practices and geographic location of the farmer, access to information on safe use of pesticide, of pesticides, and safety behavior in pesticide use. The results provide strong empirical support for the hypothesis that in reducing the effects of the pests, farmers have resorted to increasingly using more pesticides. Poor extension services, region of the farmers, acreage, frequency of pesticides application, number of crops grown, and low general knowledge on pesticide usage are attributed to overuse of different pesticides.

The smallholder vegetable production systems were informal and uncontrolled, with farmers generally lacking pest control knowledge and education on the safe use of pesticides. Agricultural land was highly fragmented with farmers operating in small pieces of land. The farming population involved was generally young with low level of education. Insecticides followed by fungicides and herbicides were the main pesticides used. These pesticides had been previously reported in Tanzania [12,14] and elsewhere [7,15,16].

The levels of pesticides volumes were considerably high in smallholder vegetable production with increased frequency application, unlike low levels of pesticides use reported in rice production [17]. Farmers sprayed on average once every week throughout the farming season. Previous studies [7,18,19] reported similar practices of overuse and abuse of pesticides by farmers. Large quantities of pesticides result in high accumulation of pesticides residues in the environment and food produce [20,21], affecting biological diversity of the ecosystem. Limited knowledge on pesticide use among farmers is assumed to be the reason for the non-use of the recommended amounts of pesticides [22]. Owing to the high volumes of pesticides sprayed, high levels of pesticides residues are presumed to be found in vegetable produce and the environment under smallholder vegetable production systems in Tanzania.

Pesticide use has increased tremendously in the recent past. Over 60 different pesticides were used in this study, as opposed to 40 different types reported previously in vegetable farming from Tanzania [12]. However, this trend was accompanied by a changing trend of applying mixed pesticide formulations. Increasing trend on pesticides use had been previously reported [22], whereby most farmers (88.8%) reported an increasing trend in pesticide use each year. Indeed, combination of different pesticides formulations was increasingly applied. Gradual change in the application of mixed active ingredients might be due to increased pest resistance to individual organophosphate, carbamates, and pyrethroids, which have been traditionally used [1]. Increased number of pesticides has a critical implication on the health of farmers, consumers of vegetable crops, and the environment where pesticides end.

Management of empty pesticides containers fosters environmental exposure. Most farmers burnt pesticides empty containers, while others buried them. Similar disposal methods and farmers reusing the empty pesticides containers for domestics purposes have been previously reported [1,19,23,24]. Generally, farmers’ awareness of disposal and management of pesticides was low. Most empty pesticides containers were left in the field during the whole farming seasons and collected/burnt during field preparation in the next farming season. The tendency of leaving pesticide leftovers and wastes in the farm or dumping directly onto the land or into water [25] increases continuous exposure risks to both humans and wildlife through dietary and environmental exposure, soil, and air [26]. Burning of empty containers directly without triple rinsing produces toxic fumes which increases risk of respiratory health problems, while burying increases risks of contaminating both surface and underground water and adverse effects to non-target organism including aquatic and terrestrial ecosystems [27].

The reported prevalence of Class II (moderately hazardous) pesticides poses high health risks due to exposure to hazardous pesticides. Profenofos, chlorpyrifos, and cypermethrin among pesticides commonly used, have been associated with decreased ecological functioning, degradation of natural vegetation, decreased number of biological species, and the depletion of fish resources [28]. These pesticides have been reported to dominate agricultural production in developing countries [7,13,29]. Pesticides use in smallholder vegetable production system is therefore unsustainably affecting the environment, necessitating further studies on environmental exposure of pesticides use. Furthermore, organophosphates and carbamate pesticides prevailing among smallholder vegetable production indicate high risks of adverse effects of pesticides exposure. This high risk poses a remarkable health concern due to pesticides neurotoxic effects as a result of the depression of acetylcholinesterase enzyme activities [30].

The key drivers of farmers’ increased use of pesticides as estimated from of the probit regression model showed that region of the farmers, number of vegetable crops grown, and mixing practices of pesticides were significantly associated with the farmers’ likelihood of using high levels of pesticides.

In regions with persistent use of pesticides, farmers are persuaded to use increased levels of pesticides continuously. Similar influence of geographical location on pesticides handling practices had been reported among vegetables farmers in Kenya [24]. Farmers’ pesticides handling is, therefore, learned over experience [7], which ultimately become a common practice and lifestyle. The lifestyle of the farming population had been associated with patterned pesticides handling practices [19]. This influences farmers to increase more pesticides in efforts to combat crop pests and diseases with the belief that the more pesticides used, the more progressive the farmer is.

Most farmers cultivated more than one vegetable crop consecutively. This trend has also been found to influence farmers’ likelihood of using more pesticides. This is fueled by the growing demand for increased productivity in a limited area of agricultural land. In efforts to control multiple pests affecting their crops, they resort in using high volumes and highly concentrated pesticides mixtures. This threatens both human and environmental health, disrupting natural pest control, and predator-prey relationship in the ecosystem. Extensive use of pesticides also results in the development and evolution of pests resistance to pesticides [31].

Mixing of pesticides during spraying was also found to be a determinant for increased pesticides use. Farmers mixing more than one pesticides had been previously reported [7]. This practice has a negative impact on safety of food produced, increases the risk due to increased toxicity of pesticides due to the additive toxicity of the resultant formulations. Mixing of more than one pesticide may also result in chemical interactions among pesticides molecules resulting in more severe effects to both farmers and consumers [32]. This practice may also results in chemical reactions of pesticides components that may change the potential of active ingredients, hence, lowering the effectiveness of the pesticides due to antagonistic effects. This may explain the considerable high proportion of farmers reporting lower effectiveness of pesticides in the control of pests.

The main reasons for mixing were minimizing spraying costs and increase the knockdown effects of pesticides. Farmers’ opinion that some of the pesticides are less effective, persuades them to use mixtures [7]. Nonetheless, farmers mixed pesticides contrary to the instructions pesticide labels, implying that farmers used pesticides without the proper guidance of agricultural experts [33]. Pesticides mixing was done mainly in tanks, which fosters farmers’ direct contact with pesticides [34]. Long term exposure to pesticide increases risk of genotoxicity and carcinogenesis [35]. Such exposure subjects the farming population to the development of numerous diseases including Non-Hodgkin’s lymphoma, multiple myeloma, pancreatic, stomach, liver, bladder and gall bladder cancer, prostate cancer, Parkinson disease and reproductive outcomes due to pesticides exposure [36,37]. These health risks depend not only on how toxic the pesticide is, but also on the level of exposure [38]. Poor pesticides handling and haphazard pesticides spraying have been associated with contamination of food produce with hazardous chemicals from pesticides [39–41].

Farmers’ perception of low effectiveness of pesticides, lack of access to information on safe pesticide use, poor use of safety gears and inability to read pesticides labels influences the level of pesticides use.

The wrong perception of the effectiveness of pesticides influences farmers’ use of high pesticide concentrations, to increase the frequency of applications and mixing of several pesticides together for better control of different pests and diseases [7,18]. Access to safe use information negatively affects farmers’ attitude resorting in increased use of pesticides. A study on farmers’ training on pesticides use and safety behavior among rural farmers in northern Greece revealed that most trained farmers showed higher levels of knowledge of pesticide use, pesticide hazard control, and safety behavior than non-trained farmers [42]. Poor pesticides handling practices among smallholder vegetable producers can, therefore, be associated with lack of training on pest control and safe use of pesticides. This is because, knowledge of pesticide risks on human health increases with formal education and training on pesticides safe use [22].

Only few farmers (11.1%) had received training on pesticides safe use. Similar findings had been reported previously in Pakistan, where 12.9% had access to information about pesticide use [7]. Very few farmers having access to pesticide safe use information got from pesticides retailers. Farmers receiving information from pesticides retailers was also reported among tobacco farmers in Greece [43]. This information is mainly oral instructions, which increases the risks of misinformation. Lack of access to pesticides safe use training may further impact negatively on the productivity and performance of vegetable subsector. Previous studies had shown handling Class II WHO pesticides and receiving advice on pesticides use from pesticide retailers increase inappropriate handling practices of pesticides significantly [24]. Poor use and handling practices observed in this study may be a result of poor information on pesticides use provided by pesticides retailers.

The importance of providing farmers with useful information on pesticides used should not be emphasized. Studies have shown that the negative effects of pesticides resulting from poor use of pesticides can be minimized by educating and training farmers on judicious use of pesticide[22]. Pesticide exposure can also be managed by educating farmers and addressing conditions forcing farmers to excessively use pesticide [44]. Improved knowledge on pesticides use can ultimately avert environmental pesticide exposure through minimizing their use and replace highly toxic pesticides with those of low toxicity [45].

These findings complement what had been reported earlier that pesticide handling is worse in developing and under-developed countries. This is because safety and regulatory guidelines on pesticides handling are hardly practiced [5] and most farmers in these countries are not adequately informed about the hazards associated with the chemicals [12,46]. Farmers are, therefore, at high risk of exposure during pesticides application due to poor monitoring and lack of well-trained experts to train and guide them on the safe use of pesticides [19,47].

Most farmers did not use personal protection equipment (PPE) when handling pesticides. The use of gloves and boots, which is the minimum PPE for most pesticide products [48] was not a common practice. Similar findings were observed among farmers in Pakistan where individuals often did not PPE when working with pesticides [22,49]. Likewise, another study in northern Greece revealed low frequency of use for gloves, goggles, face mask, coveralls, and respirator [50].

In this study, wearing of PPE was found to influence negatively the use patterns of pesticides. Farmers not using PPE were more likely to excessively use pesticides. Those wearing PPE might be more conscious on the safe use, hence, precautionary using low levels of pesticides. Poor use of PPE and high application frequencies implies increased farmers’ contact hours with pesticides. This results in exposure to pesticides during preparation and application of the pesticide spray solutions and during the cleaning-up of spraying equipment [48].This also translates in increased risk of exposure because large proportion of pesticide absorbed into the body comes from dermal exposure [51].

Evidences of pesticides intoxication exerting the strongest positive influence on PPE use had been reported [50], highlighting the importance of PPE use in reducing pesticides exposure among farmers. Increasing farmers’ awareness on the proper use of PPE would also result in the judicial use of pesticides. Less use of pesticides and the correct use of the appropriate type of personal protective equipment in all stages of pesticide handling and reduce pesticides exposure [48].

Reading of pesticides label negatively contributed to increased use of pesticides. As revealed from the findings, majority of the farmers did not read pesticides labels and consequently did not use the information provided therein. Similar findings had been reported among Greek tobacco farmers where only few farmers relied on the information found on the product label, implying that essential information about pesticide handling and safety issues found on pesticide labels are not effectively communicated to farmers [43]. As a result, farmers who did not read instructions on pesticides labels were more likely to excessively use high level of pesticides. Owing to the fact that majority of farmers had low level of education, this could imply limited reading habit which could consequently affect their ability to read and utilize useful information from pesticide labels. Similarly, in situations of farmers’ low levels of education, reading the instructions written on the pesticide containers had not been found much effective [22].

Pesticides label failing to supplement farmers’ knowledge on proper use implies that they do not provide informational needs of the farming population. The use of pesticides registered for other crops as well as in the treatment of ectoparasites and ornamental crops indicates a knowledge gap in the utilization of information provided on the pesticide label. These findings support previous studies which reported that farmers could use any pesticides product in controlling pesticides. Their desire to eliminate crop pests drives them to use any pesticide, unregistered or banned pesticides product [52,53].

## 5. Conclusion

Data on this study highlight the changing trends and determinants of increased pesticides use in smallholder vegetable production systems. Determinants of increased pesticides use and drivers of farmers’ behavior in changing pesticides use practices were assessed. Although several studies had been done on farmers’ behavior on pesticides handling, and information available may be considered sufficient, determinants and drivers of increased pesticides used in smallholder vegetable production systems in Tanzania had not been sufficiently established.

High volumes of pesticides were excessively used by smallholder vegetable producers. Their use had almost doubled during the last decade with farmers generally lacking pest control knowledge and education on the safe use of pesticides. There is ineffective control and monitoring of pesticides in smallholder vegetable production with an increasing trend of pesticides use coupled with the changing trend of the pesticides market. Combination of different pesticides formulation was increasingly applied. Mixing a wide range of pesticides was a common practice. Increased volumes of pesticides sprayed on a single spray per acre indicate high levels of pesticides residues in vegetable produce and the environment. Hence farmers and the general population are at risk of a wide range of detrimental health effects due to occupational and environmental exposure, which can result from high-level exposure from various pesticides and combined pesticides formulations.

The fate of pesticides use in smallholder vegetable production systems is therefore the culmination of serious health and environmental implications. Excessive pesticides use in controlling crop pests and diseases, escalated by increased number of crops, improper use of PPE, mixing practices of different pesticides subjects the general population to pesticides environmental exposure thereby jeopardizing sustainability of smallholder vegetable production in Tanzania. Low level knowledge, high frequency of pesticide use and mixing practices imply that both human and environmental exposure to pesticides is a serious matter of concern.

These findings signify some policy implications. Pesticides have unintended consequences regardless of the use or misuse by the smallholder vegetable farmers. Changing pesticide handling practices necessitates review of the current registration process of pesticides by considering the use of greener pesticides to minimize the effects as well as the reconsideration of the newer formulations that are safer and can replace the more toxic and highly hazardous pesticides

Poor pesticides use and associated health and environmental risks can be minimized through effective extension system to build farmers’ capacity and monitor pesticides use at farm level. The public based extension system needs to be restructured and well-coordinated targeting specific value chains, including smallholder vegetable production systems.

Regular training to extension officers on current and emerging issues of pest control safe use of pesticide is vital. Since pesticides retailers play a significant role in proving farmers with pesticides safe use information, mandatory training on safe use of pesticides and regular training is also important. The changing patterns of using combinations of different pesticides need to be studied further. Pest resistance to individual pesticides formulation needs to be scientifically established, and reason for pesticides mixing should be justified scientifically. Environmental exposure studies are also needed to establish the ecological effects of the changing patterns of pesticides use.

## Acknowledgments

The author expresses appreciation to farmers, village extension officers as well as village government in respective villages for their support and organizing the household surveys and data collection.

